# Noncanonical Wnt/Ror2 signaling regulates cell-matrix crosstalk to prompt directional tumor cell invasion and dissemination in breast cancer

**DOI:** 10.1101/2021.05.15.444295

**Authors:** Hongjiang Si, Na Zhao, Andrea Pedroza, Jeffrey M. Rosen, Chad J. Creighton, Kevin Roarty

**Affiliations:** Department of Molecular and Cellular Biology, Baylor College of Medicine, Houston, TX 77030; Dan L. Duncan Comprehensive Cancer Center, Breast Cancer Program, Baylor College of Medicine, Houston, TX 77030; Department of Medicine, Baylor College of Medicine, Houston, TX 77030

## Abstract

Bidirectional cell-extracellular matrix (ECM) interactions represent fundamental exchanges during tumor progression. We demonstrate the noncanonical Wnt receptor, Ror2, regulates tumor cell-driven matrix remodeling in models of breast cancer. Wnt/Ror2 loss-of-function triggers tumor cell invasion, accompanied by changes in actin cytoskeleton, adhesion, and collagen crosslinking gene expression programs. E-cadherin levels decline upon Ror2 depletion, and spatially, we pinpoint the upregulation and redistribution of α5 and β3 integrins together with the production of fibronectin in areas of invasion. Wnt/β-catenin-dependent and Wnt/Ror2 alternative Wnt signaling appear to regulate distinct functions for tumor cells regarding their ability to modify cell-ECM exchanges during invasion. Furthermore, blocking either integrin or focal adhesion kinase (FAK), a downstream mediator of integrin-mediated signal transduction, abrogates the enhanced migration observed upon Ror2 loss. These results reveal a critical function for the alternative Wnt receptor, Ror2, as a determinant of reciprocal communication between tumor cells and ECM during cancer invasion and metastasis.

## INTRODUCTION

Early steps of cancer metastasis require that tumor cells actively invade and disseminate from the primary tumor site by breaching the basement membrane and acquiring access to vasculature to spread to secondary organs^1^. Such a process requires that tumor cells engage signaling pathways that enable remodeling of the actin cytoskeleton while simultaneously tuning cell-cell and cell-matrix adhesions during the invasion and cellular transit within the surrounding stroma^2^. Cellular and microenvironmental heterogeneity in cancer, together with variations in migration strategies, have hindered recent efforts to thwart the initial invasion and dissemination stages responsible for spurring cancer metastasis ^3^.

Wnt signaling is a known regulator of cell fate, migration, and polarity during key morphogenic embryonic and postnatal development events. Often such patterning requires simultaneous cell fate specification concomitantly with spatial positioning of the cells to achieve coordinated morphogenesis and dynamic cell movements^4^. While canonical Wnt/ β-catenin-dependent signals stabilize intracellular β-catenin and dictate self-renewal and cell-fate choices, alternative Wnt/ β-catenin-independent cues coordinate various processes associated with cell movement, such as planar cell polarity and convergent extension^5^. Like patterned organisms and tissues, tumors comprise a hierarchy of cell types with a range of molecular and phenotypic heterogeneity^6^. Such cellular diversity is often accompanied by the presence of spatial and temporal signaling changes during tumor progression; however, it remains unclear how such evolutionarily conserved pathways regulate tumor cell behaviors during cancer progression.

ECM reorganization often accompanies tumor cell invasion to provide an appropriate scaffold for tumor cell anchorage and rewires intracellular molecular signals that instigate tumor cell behaviors during cancer progression. In breast cancer, fibrosis and ECM stiffening are associated with poor prognosis in patients^7-10^. Matrix proteins like fibronectin are assembled with type I collagen and enable lysyl oxidase (LOX)-or LOX-like dependent collagen crosslinking and organization^11^. Tumor cells respond to altered ECM composition primarily through integrin receptors, the physical link between the actin cytoskeleton within the cell and the outside ECM. Such cell-ECM interactions prompt focal adhesion kinase (FAK) autophosphorylation within the cell to reorganize its actin cytoskeleton, promote the assembly/disassembly of focal adhesions complexes, and apply traction forces to ECM proteins to facilitate directional cellular migration^12^. Like morphogenic and patterning processes in development, cancer cells undergo dynamic fluctuations in cell polarity, cytoskeletal organization, and cell-cell cohesion during cellular movement, highlighting the importance of tumor cell coordination and reciprocal engagement with their surrounding microenvironment^13,14^. However, the precise regulation of such cellular behaviors and the signals that trigger invasive cell behavior in the context of cancer progression remain unknown.

We previously discovered that the intratumor landscape within basal-like Triple-Negative Breast Cancer (TNBC) harbor tumor cells that differentially utilized distinct modes of Wnt signaling, demarcated by canonical Wnt/β-catenin-active cells co-existing with tumor cells expressing Ror2, an alternative β-catenin-independent Wnt receptor^15^. Here, we identify that genetic depletion of Ror2 in primary tumors yields increased stromal remodeling and collagen deposition, a poor prognostic factor for breast cancer patients^16-18^. In the present study, we use a 3-dimensional (3D) Type 1 collagen model of tumor organoids to detect gene expression programs representing cell adhesion, collagen fibril organization, and cell-ECM alterations resulting from Ror2 loss. Junctional E-cadherin expression is compromised upon Ror2 depletion, accompanied by increased tumor cell invasion into the surrounding collagen matrix. In addition, we uncover the upregulation of integrin-mediated signaling in Ror2-deficient tumor cells, particularly the expression integrin-α_5_ integrin and integrin-β_3_ at the invasive front of tumor organoids. Intriguingly, fibronectin (FN) is concomitantly upregulated at sites of invasion upon Ror2-loss. Such changes in integrin receptor abundance and localization trigger changes in the cytoskeletal architecture of Ror2-deficient tumor cells, prompting an increase in F/G actin ratios and the activation of FAK. Inhibition of either integrin or FAK activation abrogates the increased invasion driven by Ror2-loss. Such changes are distinct from processes regulated by the activation of Wnt/ -catenin and offer insights into how canonical and alternative Wnt pathways cooperatively regulate the coordination of tumor cell behaviors during breast cancer progression.

## RESULTS

### Ror2 presence dictates tumor cell-directed collagen remodeling and transit through the ECM

We previously discovered the negative correlation between Wnt/β-catenin and Wnt/Ror2 alternative signaling across breast cancer subtypes and the coexistence of spatially distinct canonical Wnt/β-catenin active and Ror2-expressing β-catenin-independent tumor subpopulations within the cellular landscape of TP53-null TNBC basal-like models of breast cancer^15^. The syngeneic TP53-null transplantable tumor model is a unique library of tumors representing human breast cancer subtypes at the molecular level^19-21^. TP53-mutated (typically missense and loss-of-function) breast tumors make up 90% of TNBC, so the current models are clinically relevant^22,23^. Although Wnt pathways are evolutionarily conserved signals that are essential for the development of multicellular organisms, how distinct but interconnected Wnt pathways regulate complex tumor cell behaviors during cancer progression remains unclear. In experimental models, while Ror2 status impacts the proportion of molecularly distinct basal-like (BL) CD24^+^CD29^+^ and claudin-low (CL) CD24^-/low^CD29^+^ cells within tumors, we discovered that along with the regulation of subpopulation plasticity, tumors deficient in Ror2 had a propensity to exhibit a highly disorganized tumor stroma. We, therefore, investigated the functional implications of this altered stroma and how Wnt/Ror2 functions specifically contribute to the microenvironmental alterations evident within TNBC models of breast cancer. Trichrome staining of Ror2-depleted tumors revealed a significant increase in collagen abundance and disorganized tumor architecture relative to Ror2-intact tumors (**Figure 1A, B**). Notably, a 2-fold higher integrative density for collagen presence was determined across Ror2-depleted shRor2 tumors relative to shLUC control tumors (**Figure 1C**). Using a collagen hybridizing peptide (CHP) which specifically binds to denatured collagen, we also established that unfolded collagen, likely the result of proteolytic remodeling by tumor cells, was expanded within tumors following Ror2 loss (**Figure 1D**). In shLUC control tumors, CHP presence was predominantly localized within the adjacent stroma surrounding tumor cells, while in shRor2 tumors, CHP was expanded in adjacent stromal areas, in addition to intercellular regions between tumor cells (**Figure 1D, arrows**).

**Figure 1.**
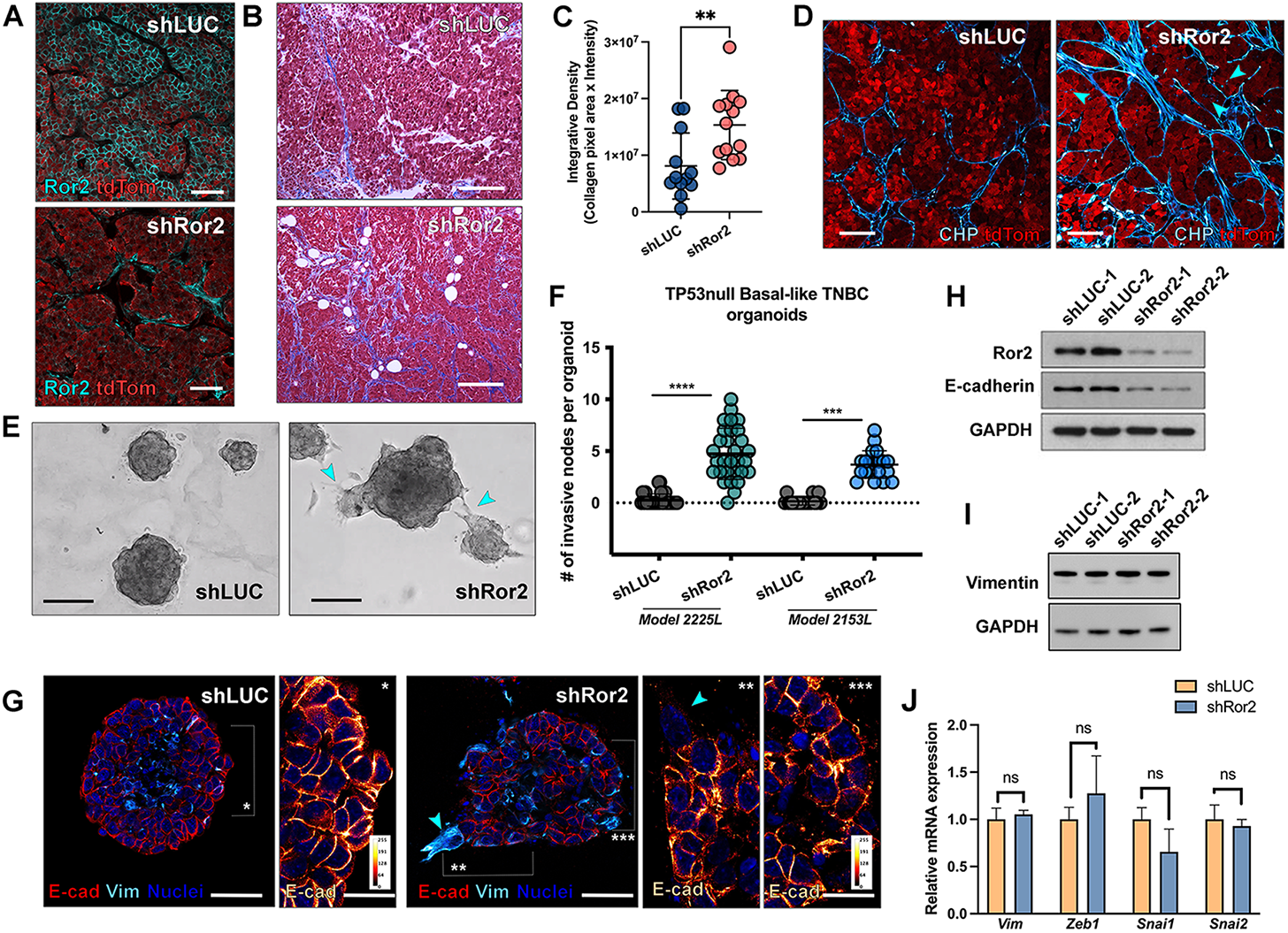
Wnt/Ror2 regulates tumor cell-directed collagen remodeling and invasion. (A) Immunofluorescence for Ror2 protein in TP53null 2225L basal-like GEM models (Cyan – Ror2, tdTomato – LeGO-hairpin transduced tumor cells). (B) Representative trichrome staining of basal-like 2225L TP53null mammary tumors showing collagen abundance (blue) and tumor organization in shLUC and shRor2 tumor sections. Scale 50 μm. (C) Quantification of Trichrome collagen abundance by Integrative Density (collagen pixel area x fluorescence intensity, n=4 tumors per group, 4 fields per tumor, ^**^p<.01). (D) Collagen Hybridizing Peptide histopathology (CHP-Alexa488, pseudocolored Cyan hot LUT) within shLUC and shRor2 2225L tumors (tdTomato – tumor cells). (E) Brightfield DIC images of shLUC and shRor2 2225L tumor organoids in Type I collagen. (Scale 50 μm). Ror2-depleted organoids exhibit increased invasive projections. (F) Quantitation of the extent of invasion in Ror2-depleted organoids within the basal-like 2225L and 2153L TP53null models. Representative quantitation of 3 independent experiments (n=30 organoids per group, ^****^p<.0001, ^***^p<.001). (G) Immunofluorescence of E-cadherin (red), vimentin (cyan), and nuclei (blue) in 2225L shLUC and shRor2 organoids, showing downregulated E-cadherin and altered distribution of vimentin upon Ror2 loss. G^*^ shows E-cadherin continuity within shLUC organoids. G^**, ***^ shows discontinuity and downregulation of E-cadherin in shRor2 organoids Scale 50 μm, ^*, **, ***^ Insets 25 μm. Arrows denote areas of invasion with most downregulated levels of E-cadherin and redistributed Vimentin. (H) Western blot of 2225L shLUC and shRor2 organoids for Ror2 and E-cadherin, revealing decreased Ror2 and E-cadherin expression levels in shRor2 organoids. **(I)** Western blot of 2225L shLUC and shRor2 organoids for vimentin. (J) Quantitative SYBR Green RT-qPCR measurement of mesenchymal markers (*Vim, Zeb1, Snai1*, and *Snai2*) in 2225L shLUC vs. shRor2 tumor cells from organoid cultures. Gene expression levels of the shRor2 group was represented relative to the control shLUC group and fold changes were plotted (ns: not significant; n=3 biological replicates for each group).

To better understand the functional consequences of Wnt/Ror2 signaling on tumor cell/matrix interactions, we established 3D tumor organoid cultures within embedded Type I collagen ECM to evaluate the impact of Ror2-depletion on cell-ECM exchanges during cancer invasion. Collagen I is the most abundant scaffolding protein present in tissues, and its crosslinking is highly associated with breast cancer risk^9^. Given the altered collagen abundance in primary tumors observed upon Ror2 depletion, we reasoned that Wnt/Ror2 signaling could provide tumor cell intrinsic instruction by dictating pericellular ECM composition, and consequently, bidirectional communication between tumor cells and ECM. Accordingly, we generated tumor organoids using two independent Genetically Engineered Mouse (GEM) TP53-null transplant models^15,21^ representative of the basal-like subtype of TNBC (2225L and 2153L) to probe the functional implications of Wnt/Ror2 biology on cell-matrix interactions during tumor progression. Notably, Ror2 knockdown by lentiviral shRNA prompted the active invasion of tumor cells into the surrounding collagen-rich matrix, accompanied by a 4.5-fold and 3.7-fold increase in protrusive tumor cell extensions within 2225L- and 2153L-shRor2 organoids, respectively, as compared to non-invading control shLUC organoids (**Figure 1E, F**).

We further investigated defects in epithelial and mesenchymal states, given our previous observations that Ror2 loss prompted an enriched CL subpopulation and CL molecular signature in tumors^15^. Such states can dictate both single and collective behaviors of tumor cells during invasion and dissemination, so we reasoned that EMT could be involved in Ror2 loss-of-function (LOF). While Ror2 depletion prompted the downregulation of E-cadherin (**Figure 1H**), the mesenchymal marker, vimentin, was equally expressed between experimental and control groups (**Figure 1I**). Interestingly, shRor2 organoids harbored a disrupted pattern of E-cadherin at the cell cortex within the organoid body relative to Ror2-intact organoids by immunostaining (**Figure 1G**). This expression of E-cadherin consistently decreased as cells invaded into the surrounding ECM upon Ror2 loss (**Figure 1G**, ^**\***, **\*\***, **\*\*\***^ **insets**). Organoids did not exhibit a change in total vimentin expression levels, yet localization of vimentin shifted towards areas of active invasion in Ror2-depleted organoids at protrusive areas of ECM involvement and E-cadherin loss (**Figure 1G;** ^**\***, **\*\***, **\*\*\***^ **inset arrows**). No notable changes in other mesenchymal-associated transcription factors such as *Zeb1, Snai1*, and *Snai2* were detected upon Wnt/Ror2 signaling impairment (**Figure 1J**). Thus, the disruption in E-cadherin expression and lack of change in mesenchymal markers suggest that cell-cell adhesion is compromised upon Ror2 depletion rather than a complete shift from an epithelial to a mesenchymal state.

### Ror2 depletion prompts alterations in actin cytoskeleton and cell-ECM dynamics

The enhanced degree of invasion in tumor organoids and increased expression of collagen within Ror2-depleted tumors prompted the investigation of the molecular determinants that mediate the invasive switch downstream of Ror2. Accordingly, we performed RNA sequencing on 4-day cultures of 3D shLUC vs. shRor2 tumor organoids within Type I Collagen, derived from both basal-like 2225L and 2153L TP53-null GEM models (**Figure 2A, Figure S1A**). Gene ontology programs encompassing the guidance of tumor cell movement, particularly the regulation of cell migration, cell-cell adhesion, integrin-mediated adhesion, collagen fibril organization, and cytoskeletal organization, were highly represented upon *Ror2* knockdown (**Figure 2B-D, Figure S1B**). Given the manifestation of collagen fibril expansion upon Ror2 loss, we next investigated how Ror2 presence might shape states of intercellular cohesion and reciprocal exchanges between tumor cells and the ECM. Interestingly, Lox and Lox-like 2 (Loxl2), enzymes required for the biogenesis and crosslinking of fibrillar collagen, were significantly upregulated upon Ror2 loss at the RNA and protein levels (**Figure 2C-D**), along with tumor cell intrinsic collagen expression (**Figure 2B-D, Figure 1B-D**). Interestingly, α_5_ integrin and β_3_ integrin, receptors for ECM components responsible for physically bridging internal and external filamentous networks to enable bi-directional transmission of signals across the plasma membrane, also were increased in tumor cells following Ror2 depletion (**Figure 2D-F, Figure S1C**). From these data, we postulated that Wnt/Ror2 signaling regulates a critical signaling juncture for tumor cells by shaping integrin presence, tumor cell engagement with the ECM, and downstream cytoskeletal changes conducive for tumor cell movement.

**Figure 2.**
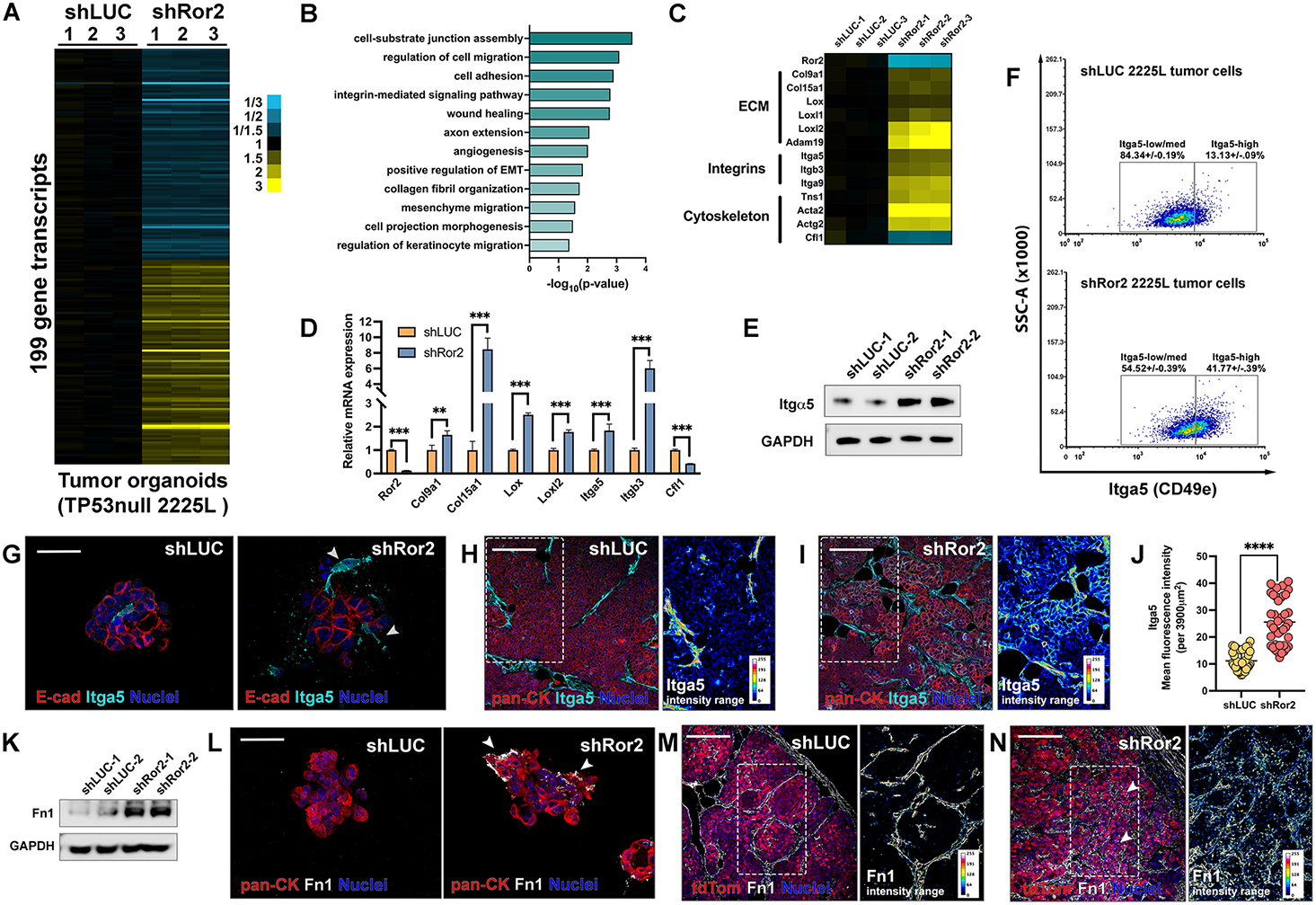
Ror2 depletion promotes gene expression alterations related to cell-ECM interactions, reshaping integrin signaling and ECM composition. (A) Heatmap display of significantly differentially expressed genes (p<0.01, t-test using log2-transformed values and fold change>1.2) in 2225L shLUC and shRor2 organoids by RNA sequencing. Fold changes are represented by two-way gradients to blue (downregulation) and yellow (upregulation). (B) Gene ontology analysis by DAVID Bioinformatics Database demonstrating the enrichment of gene expression in several biological processes upon Ror2 loss. (C) Heatmap display of highlighted genes in 2225L shLUC and shRor2 organoids related to cell-ECM interaction, integrin signaling, and cytoskeleton reorganization. Fold changes are represented by two-way gradients to blue (downregulation) and yellow (upregulation). (D) Quantitative RT-qPCR measurement of highlighted genes in 2225L shLUC and shRor2 cells for representative genes from (C). Gene expression levels were normalized to GAPDH and the shRor2 group was represented relative to the shLUC group. Fold changes were graphed (^***^: p<0.0001; ^**^: p<0.001; n=3 for each group). (E) Western blot of 2225L shLUC and shRor2 organoids for Itga5, showing increased Itga5 expression levels in Ror2-depleted organoids. (F) Flow cytometry analysis for α5 integrin Itga5 (CD49e) on shLUC and shRor2 tumor cells from the 2225L TP53null GEM model. shRor2 tumors exhibit an increase in Itga5 expression with ∼3-fold increase in Itga5-high cells upon Ror2 loss (10,000 single-cell events analyzed per tumor; plots represent n=3 tumors per group; representative of 3 independent experiments). (G) Immunofluorescence of integrin α5 (cyan), E-cadherin (red), and nuclei (blue) in 2225L shLUC and shRor2 organoids, showing upregulated integrin α5 upon Ror2 loss, particularly in invading projections (arrows). Scale 40 μm. (H) Immunofluorescence of α5 integrin (cyan), pan-Keratin (red), and nuclei (blue) in 2225L shLUC primary tumor sections, showing topographic distribution of α5 integrin in control tumors. Inset represents 16-color rainbow LUT distribution of Itga5 presence based on a linear gradient (0 to 255) in the tumor. Scale 100 μm. (I) Representative immunofluorescence of α5 integrin (cyan), pan-Keratin (red), and nuclei (blue) in 2225L shRor2 primary tumor sections, showing enhanced α5 integrin expression within and around tumor cells in shRor2 tumors. Scale 100 μm. Inset represents 16-color rainbow LUT distribution of Itga5 presence based on a linear gradient (0 to 255) within the tumor. Scale 100 μm. (J) Quantitation of Itga5 mean fluorescence intensity in (H,I) per 390μm^2^ area (n=6 tumors per shLUC and shRor2 group (^****^: p<0.00001). (K) Western blot of 2225L shLUC and shRor2 organoids for fibronectin 1, showing increased fibronectin 1 (Fn1) expression levels in Ror2 depleted organoids. (L) Immunofluorescence of fibronectin 1 (white), pan-Keratin (red), and nuclei (blue) in 2225L shLUC organoids and 2225L shRor2 organoids, showing increased fibronectin 1 deposition upon Ror2 loss, specifically around invasive projections at the tumor cell-matrix interface. Scale 40 μm. (M) Immunofluorescence of fibronectin 1 (white), tdTomato tumor cells (red), and nuclei (blue) in 2225L shLUC primary tumor sections, showing deposition of fibronectin 1 predominantly within adjacent stromal regions surrounding tumor cells. Scale 100 μm. (N) Immunofluorescence of fibronectin 1 (white), tdTomato tumor cells (red), and nuclei (blue) in 2225L shRor2 primary tumor sections, showing increased tumor cell-intrinsic fibronectin 1 deposition in shRor2 tumors along with adjacent stromal Fn1 presence. Scale 100 μm. Insets in M and N represent LUT 16-color rainbow representation of Fn1 expression based on a linear gradient (0 to 255) within the tumor (boxed areas of interest).

### Loss of Ror2 triggers the upregulation and redistribution of integrins to assemble fibronectin at sites of invasion

Integrins are transmembrane receptors that mediate cell adhesion, migration, and enable the bidirectional signal transduction between cells and ECM proteins^24^. Given the expansion of collagen presence *in vivo* within Ror2-deficient tumors, as well as the induction of collagen crosslinking enzymes Lox and Loxl2 in Ror2-depleted organoids (**Figure 2C**,**D**), we next examined the integrin changes prompted by Ror2 loss based on RNA sequencing, given their involvement in both the biogenesis of and interaction with ECM proteins during several cell migration processes^25^. In shRor2 organoids, we determined that α_5_ integrin was upregulated at both RNA and protein levels relative to control shLUC organoids (**Figure 2C-F**). We additionally detected an increase in β_3_ integrin by flow cytometry upon Ror2 loss (**Figure S1C**). Interestingly, the analysis of α_5_ integrin expression by immunofluorescence in tumor organoids revealed both upregulation and spatial repositioning within Ror2-deficient organoids, specifically at sites of tumor cell invasion within the surrounding Type I collagen matrix (**Figure 2G**). These data suggest that loss of Wnt/Ror2 signaling in TNBC tumor cells potentially primes invasion by adjusting cell-cell and cell-ECM signaling.

Integrins are α/β heterodimeric transmembrane receptors, and when the α_5_ subunit of integrin associates with β_1_ integrin, this complex forms a fibronectin receptor^25,26^. Since tumor organoid cultures were grown in Type I collagen, we investigated whether tumor-cell derived fibronectin expression was induced following Ror2 depletion. Integrins play a central role in fibronectin matrix assembly and have a requisite role in cell adhesion, invasion, and bidirectional signaling. Moreover, the deposition and restructuring of collagen in the ECM depends on the presence and stability of fibronectin. Indeed, we discovered that fibronectin protein levels were elevated following Ror2 loss in tumor organoids. This increase was most likely at the post-transcriptional level, since we did not detect induction of fibronectin mRNA (**Figure 2A,K**). Moreover, like α_5_ integrin, fibronectin was upregulated and spatially repositioned adjacent to the invading cells disseminating from the organoid body upon Ror2 loss, particularly at the sites of cell protrusions into the ECM (**Figure 2L**). Intriguingly, the *in vivo* intratumoral topology of both α_5_ integrin and fibronectin expanded from the tumor periphery in controls to a more sizeable area encompassing both the periphery and inner tumor mass following Ror2 loss (**Figure 2H-J, M-N**). Though integrin and fibronectin presence were noted in the adjacent stromal regions within both shLUC and shRor2 tumors, tumor cell intrinsic expression was uniquely apparent in Ror2-deficient tumors based on an increase in mean fluorescence intensity within such regions (**Figure 2J, M-N**). Collectively, these data suggest that tumor cell-derived fibronectin, dictated by Ror2 expression, helps facilitate integrin-mediated invasion into the adjacent cellular matrix.

### Focal adhesions and actin cytoskeleton dynamics are regulated by Ror2

Integrin-linked focal adhesion complexes serve as a physical scaffold for the cell to adhere to the exterior ECM and additionally interact with the interior actin cytoskeleton, providing instructive and highly tunable cues necessary for modulating adhesion and passage through the ECM^12^. Focal adhesion kinase (FAK) is activated downstream of integrins, typically in response to integrin clustering. Given the upregulation and localization of α_5_ integrin at plasma membrane sights of invasion in Ror2-deficient tumor organoids, we suspected that this might be accompanied by the activation of FAK in both organoids and tumors with either intact-or deficient-Ror2 status. Accordingly, the level of p-FAK was upregulated in Ror2-deficient tumor organoids (**Figure 3A**). This observation was confirmed by the analysis of p-FAK immunofluorescence in 3D tumor organoids (**Figure 3E**). When probing the intratumoral landscape of p-FAK *in vivo*, we observed the prevalence of p-FAK within tumor cells located within the tumor periphery, while p-FAK levels were negligible within the tumor body central to the tumor margin in controls (**Figure 3B, C**). Interestingly, Ror2-depleted tumors had an overall increase in p-FAK activation throughout the tumor, in contrast to negligible levels of p-FAK within the body of control tumors (distal to the tumor periphery), and p-FAK presence was significantly expanded following Ror2 loss (**Figure 3C**). These results reveal the significant change in focal adhesion topology dictated by the presence of Wnt/Ror2 signaling *in vivo* (**Figure 3B-E**).

**Figure 3.**
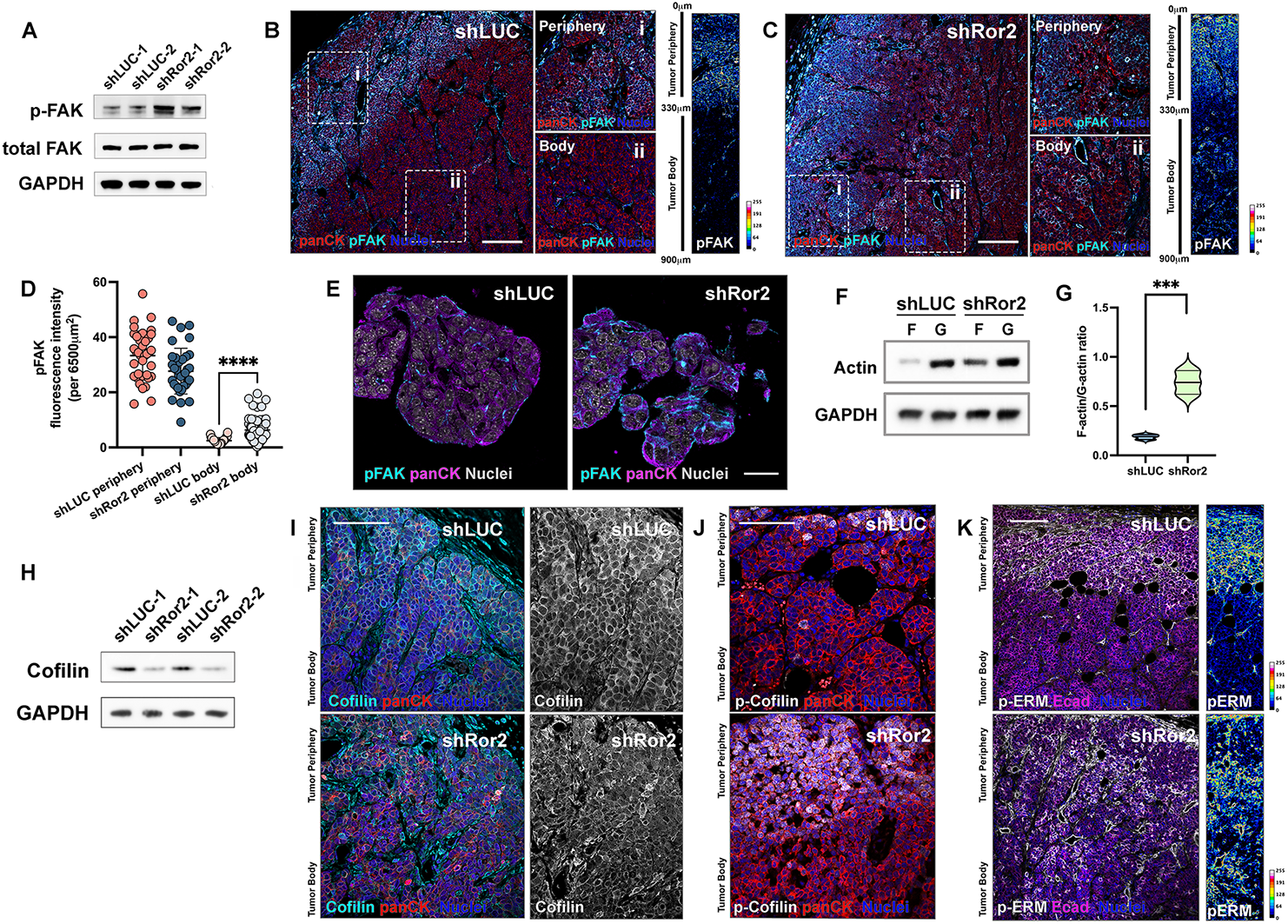
Focal adhesions and actin cytoskeleton dynamics are regulated by Ror2. (A) Western blot of 2225L shLUC and shRor2 organoids for phosphorylated FAK (Y861) and total FAK showing increased FAK phosphorylation in Ror2 depleted organoids. (B) Immunofluorescence of p-FAK (cyan), pan-CK (red), and nuclei (blue) in 2225L shLUC primary tumor sections, showing topographic distribution of p-FAK in control tumors. Scale 150 μm. (B^i^) Prevalent p-FAK staining in the periphery of shLUC tumors. (B^ii^) Limited p-FAK staining in the body of shLUC tumors. Scale 100 μm. (C) Immunofluorescence of p-FAK (cyan), pan-CK (red), and nuclei (blue) in 2225L shRor2 primary tumor sections, showing topographic alterations in p-FAK distribution upon Ror2 loss. Scale 100 μm. (C^i^) Prevalent p-FAK staining in the periphery of shRor2 tumors. (C^ii^) Expansion of p-FAK staining in the body of shRor2 tumors. Scale 100 μm. (D) Quantitation of pFAK immunofluorescence intensity per 6500 m^2^ area, distinguishing between periphery and inner tumor body of shLUC and shRor2 tumors. Statistically significant increase in pFAK fluorescence intensity is shown (^****^p<.00001, n=3-4 tumors per group, 10 regions within periphery and body per tumor evaluated). (E) Immunofluorescence of p-FAK (cyan), pan-CK (magenta), and nuclei (grayscale) in 2225L organoids, showing changes in p-FAK levels and distribution within protrusions upon Ror2 loss. Scale 25 μm. (F) F-actin/G-actin ratio assay of 2225L shLUC and shRor2 cells showing increased F-actin content in Ror2 depleted cells. (G) Quantification of the F-actin/G-actin ratio in 2225L shLUC and shRor2 cells (^***^: p<0.0001; n=3 biological replicates). (H) Western blot of 2225L shLUC and shRor2 organoids for cofilin showing reduced cofilin expression in Ror2 depleted organoids. (I) Immunofluorescence of cofilin (cyan), pan-CK (red), and nuclei (blue) in 2225L shLUC and shRor2 primary tumor sections, showing expression and localization of cofilin. Grayscale (right) represents cofilin cytoplasmic staining in shLUC control tumors and diminished and irregular staining in shRor2 tumors. Scale 75 μm. (I′) Cofilin staining in the periphery of shLUC tumor. (J) Immunofluorescence of p-cofilin (white), pan-CK (red), and nuclei (blue) in 2225L shRor2 primary tumor sections, showing increased p-cofilin Ser3 upon Ror2 depletion, particularly more within the tumor body. Scale 75 μm. (K) Immunofluorescence of p-ERM (white), pan-CK (magenta), and nuclei (blue) in 2225L shLUC and shRor2 tumors. Loss of Ror2 expands pERM localization throughout the tumor. Scale 100 μm. Right insets show color coded increased and redistributed p-ERM upon Ror2 depletion by LUT 16-color linear gradient (0-255) within periphery and tumor body of shLUC and shRor2 groups. Scale 100 μm.

Because of altered p-FAK activation downstream of Ror2, we suspected that actin cytoskeletal dynamics are perturbed, as indicated by RNA sequencing analysis (**Figure 2A**). Actin cytoskeletal remodeling is a cyclical process necessary for locomotion and shape demands of the cell, requiring actin binding proteins (ABPs) and fluctuations in globular monomeric (G) and filamentous (F) actin. Indeed, cytoskeletal rearrangements were identified based on the evaluation of F-actin/G-actin turnover **(Figure 3F, G)**. Ror2-depleted tumor cells exhibited an almost 4-fold higher level of filamentous (F) to monomeric (G) actin relative to Ror2-intact tumor cells (**Figure 3F, G**). Such findings support that Wnt/Ror2 signals normally maintain an actin cytoskeletal homeostasis within tumor cells by maintaining appropriate cell-cell and cell-ECM cues. Accordingly, cofilin, a key actin-binding protein responsible for the depolymerization of filamentous F-actin^27^, was significantly downregulated in tumor organoids and *in vivo* upon Ror2 loss (**Figure 3H-J**). However, cofilin downregulation was not a consequence of Wnt/β-catenin activation in the absence of Wnt/Ror2 signaling (**Figure 4I, Figure S2A, C)**). Additionally, we detected an increase in the extent of p-Ser3 Cofilin at the periphery of shRor2 tumors relative to a very narrow zone directly abutting the tumor-stromal interface in shLUC control tumors (**Figure 3J**). Ser3 phosphorylation inactivates cofilin, suggesting that heightened p-cofilin levels following Ror2 loss promote actin filament elongation, an increase in F/G actin ratio, and consequently the invasive potential of tumor cells. Lastly, Ezrin, Radixin, and Moesin (ERM), linkers of the actin cytoskeleton to the plasma membrane, were phosphorylated/activated at the tumor periphery of control shLUC tumors but were expanded spatially throughout the tumor body upon Ror2 loss (**Figure 3K**). Moreover, while pERM was located at the cell cortex of shLUC tumor cells, a discontinuous punctate localization of pERM was prevalent in shRor2 tumor cells, likely reflecting altered intercellular and cell-ECM interactions. These data collectively demonstrate a pivotal role for Wnt/Ror2 status in dictating pericellular ECM composition and integrin signaling, the outcome of which shapes the formation of focal adhesions, cellular actin cytoskeletal changes, and the transmission of ECM-derived signals necessary to move the tumor cell body.

**Figure 4.**
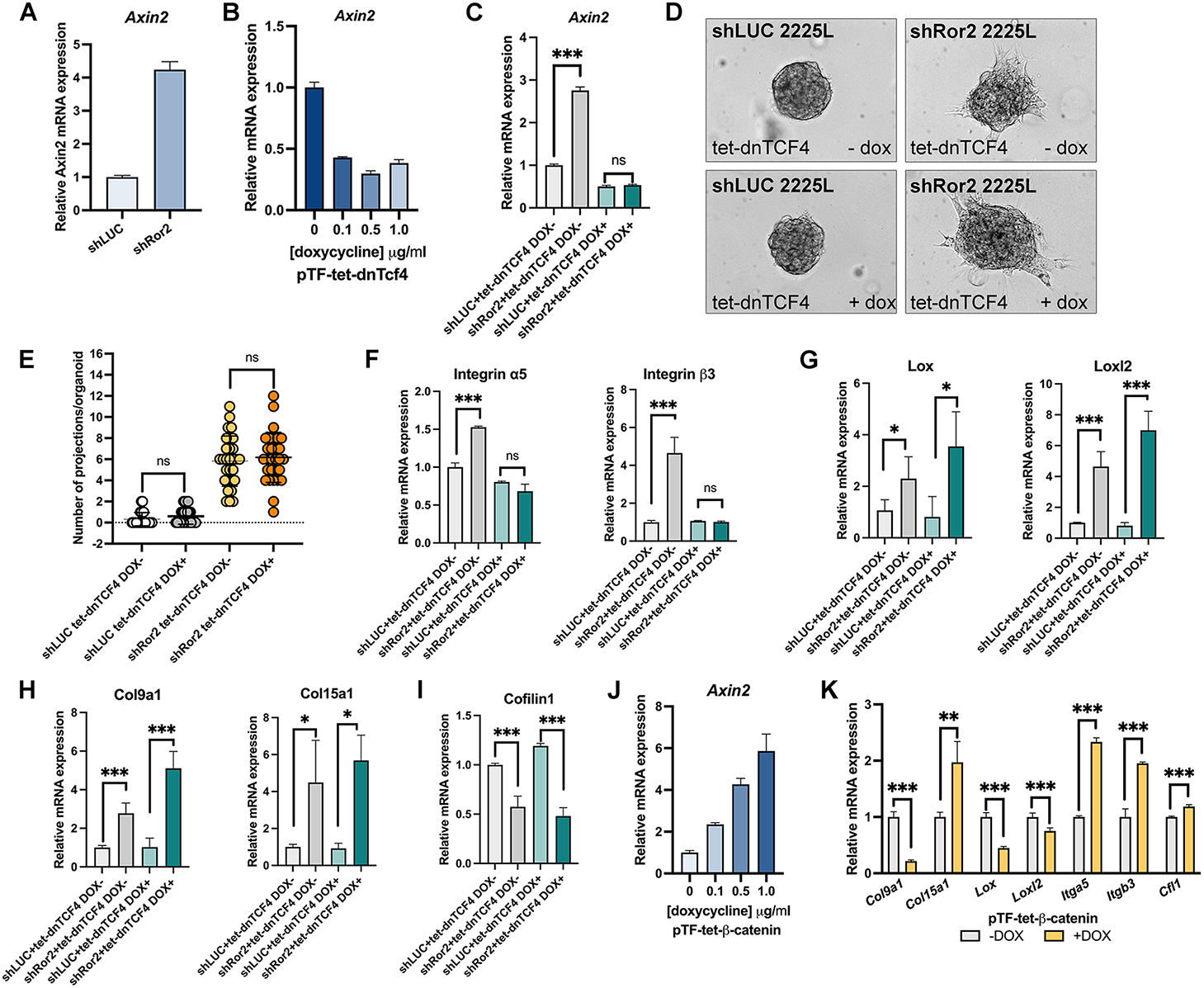
Wnt/Ror2 and Wnt/β-catenin pathways differentially regulate ECM composition and cell-ECM interactions during tumor cell invasion. (A) RT-qPCR for *Axin2* in shLUC vs. shRor2 organoids demonstrating that Wnt/β-catenin signaling is induced upon Ror2 depletion in 2225L tumor cells. (B) RT-qPCR for *Axin2* after 72 h post doxycycline induction of a pTF-tet-dnTcf4 lentiviral construct in 2225L tumor cells to block Wnt/β-catenin signaling. Shown is a dose-dependent inhibition of *Axin2* upon dnTcf4 induction. (C) RT-qPCR for *Axin2* after 72 h post doxycycline induction of a pTF-tet-dnTcf4 lentiviral construct in shLUC and shRor2 2225L tumor cells showing the inhibition of *Axin 2* expression upon Ror2 loss and dnTcf4 induction. (D) Brightfield DIC images of shLUC and shRor2 organoids in the presence or absence of dnTcf4 induction. Shown are representative images after 3 days of culture post dnTcf4 induction. (E) Quantitation of invasion in shRor2 vs. shLUC organoids after blocking Wnt/β-catenin signaling by dnTcf4. (n=30 organoids per group, representing 3 independent experiments). (F-I) RT-qPCR for ECM-associated genes in shLUC vs. shRor2 tumor cells upon dnTcf4 induction, including (F) *α5 integrin* and *β3 integrin* (G) *Lox* and *Loxl2* (H) *Col9a1* and *Col15a1* (J) *Cofilin 1*. (J, K) Inducible expression of -catenin within 2225L tumor cells and RT-qPCR for (J) *Axin2* induction following dose-dependent administration of doxycycline for 72 h and (K) RT-qPCR of ECM-associated genes *Col9a1, Col15a1, Lox, Loxl2, Itga5, Itgb3*, and *Cfl1* showing different outcomes following - catenin induction compared with Wnt/Ror2 impairement.

### Wnt/Ror2 and Wnt/β-catenin pathways differentially regulate tumor cell-ECM exchanges during invasion

Wnt/Ror2 signaling can repress Wnt/β-catenin signaling depending on the cellular context. The spatial segregation of Wnt/β-catenin active vs. Wnt/Ror2 subpopulations *in vivo* and our observation that Wnt/Ror2 can repress Wnt/β-catenin activity prompted us to assess the contribution of heightened Wnt/β-catenin activity observed in the absence of Ror2^15,28,29^. We confirmed an increase in Wnt/β-catenin signaling activity, assessed by the induction of the downstream negative feedback regulator *Axin2*, upon Ror2 knockdown within TP53null 2225L tumor cells (**Figure 4A**). We next tested Wnt/β- catenin loss-of-function in the context of Ror2 depletion by conditionally overexpressing a dominant negative form of the TCF4 transcription factor in shLUC and shRor2 organoids. We confirmed that induction of dnTCF4 reduced *Axin2* expression in a dose-dependent manner, and moreover, suppressed the elevated Wnt/β-catenin-dependent *Axin2* expression brought about by Ror2 depletion (**Figure 4B, C**). We next determined the contribution of Wnt activation on the invasion of tumor cells. Interestingly, Wnt/β-catenin LOF did not impede invasion prompted by Ror2 knockdown, suggesting the dissemination of tumor cells results from impaired Wnt/Ror2 signaling and not the ectopic activation of Wnt/β -catenin signaling in this context (**Figure 4D, E**). We additionally assessed ECM-associated gene expression changes which occurred upon Wnt/β -catenin LOF in shLUC and shRor2 organoids. We determined that Wnt/β-catenin LOF impaired integrin upregulation (*Itga5, Itgb3*) upon Ror2 loss, but was dispensable for the deposition of tumor cell-intrinsic ECM (*Col9a1, Col15a1*) and expression of Lox family enzymes downstream of Wnt/Ror2 signal loss (**Figure 4F-H**). Moreover, inducible activation of Wnt/β-catenin alone in tumor cells elicited distinct effects as compared to Wnt/Ror2 on EMT- and ECM-associated gene networks (**Figure 4J, K, Figure S2B-D**), where Wnt/β-catenin activation prompts the upregulation of mesenchymal genes despite the inability to alter ECM composition. These data suggest that Ror2 LOF phenotypes in this context are not a consequence of ectopic Wnt/β-catenin activation^15^. Thus, canonical Wnt/β-catenin-dependent signaling appears to function differently in shaping cell-ECM exchanges and invasion as compared with Wnt/Ror2-specific roles. Though distinct at the molecular level, the balance of Wnt/Ror2 and Wnt/β-catenin signaling outputs clearly impact both intercellular crosstalk and cell-ECM behavior to shape adhesion and invasion dynamics where such pathways are likely operating in concert within a tumor.

### Inhibition of integrin and FAK impedes directional migration prompted by Ror2 loss

To determine the functional implications of integrin and FAK activation on Ror2 loss-of-function phenotypes with respect to invasion and dissemination, we tested small molecular inhibitors of both integrin and FAK activation. ATN-16, a novel small molecule peptide antagonist of α_5_β1 integrin, successfully inhibited pFAK activation induced by Ror2 loss in a dose-dependent manner (**Figure 5A**). We subjected 2225L TP53null 2D cells to an in vitro wound healing assay to test the outcome of ATN-16 on the ability of shRor2 cells to efficiently close the wound. After 16 hours, shRor2 cells were more efficient at filling the area of wounding than control shLUC cells; however, ATN-161(10µM) resulted in a significantly greater impairment of Ror2-deficient cell migration and wound closure than shLUC control cells (**Figure 5B-D**). The inhibitor PF-562271 Besylate, a potent ATP-competitive reversible inhibitor of FAK activation, was also tested given the elevated p-FAK levels in both *in vitro* organoids and *in vivo* tumors upon Ror2 loss. Treatment with besylate successfully mitigated the activation of FAK phosphorylation upon Ror2 depletion (**Figure 5E**). Moreover, migration of shRor2 tumor cells was greatly reduced upon besylate treatment in the 2D would healing assays (**Figure 5F, G**). Such a reduction in migration and invasion was additionally observed in 3D organoid cultures which represent a more physiologically relevant cell-ECM landscape (**Figure 5H-J**). Thus, these studies place integrin-mediated signaling and FAK activation downstream of Wnt/Ror2 signaling. These findings have important implications for the regulation of tumor cell-ECM behaviors downstream of Wnt/Ror2 signaling governing decisions of cell adhesion, invasion, and survival during breast cancer metastasis.

**Figure 5.**
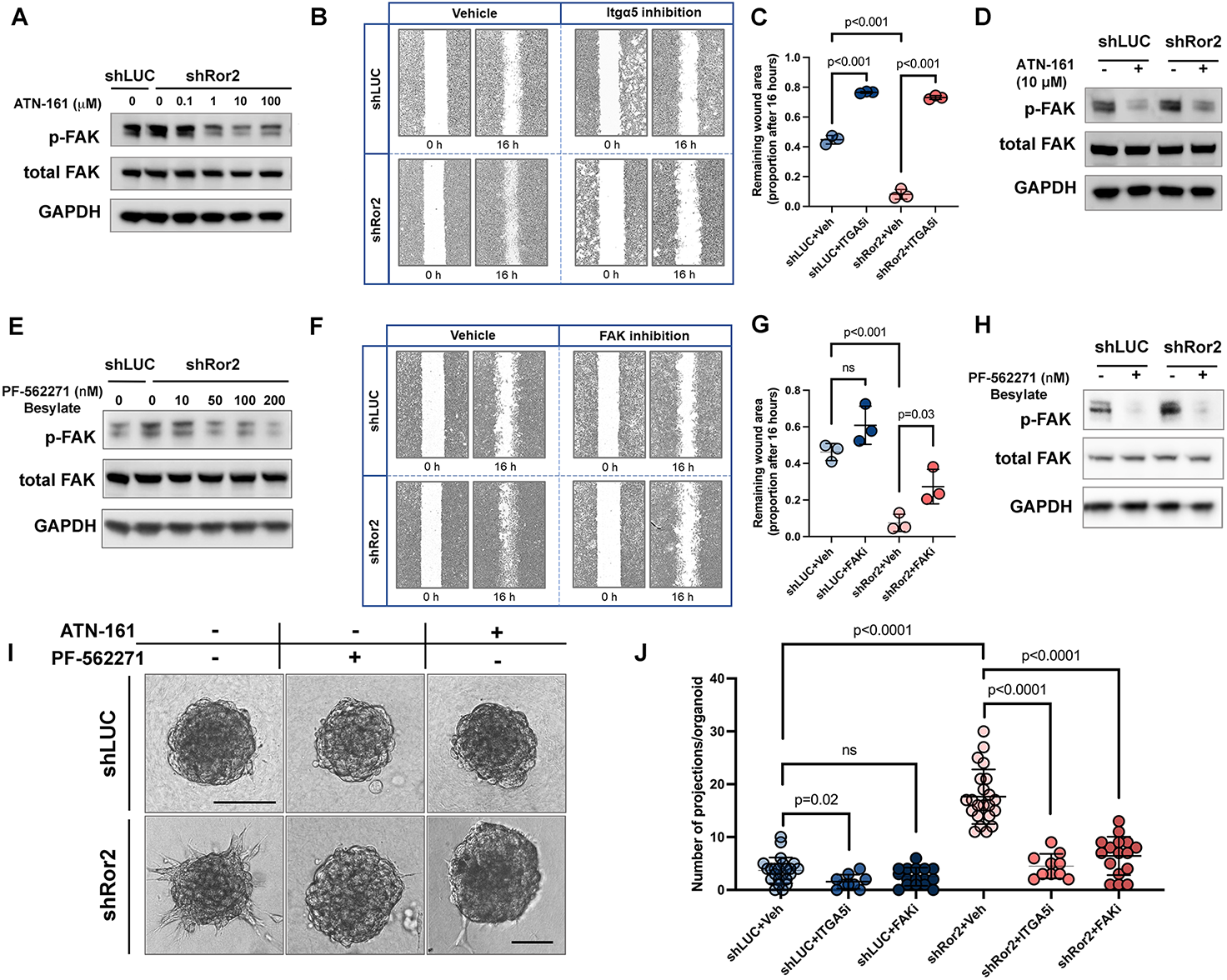
Inhibition of α5 integrin or FAK inhibit breast cancer cell migration and invasion downstream of Ror2 loss. (A) Western blot of 2225L shLUC and shRor2 cells for phosphorylated FAK and total FAK after 0, 0.1, 1, 10, or 100 μM ATN-191 treatment. (B) Brightfield images of wound healing assays on 2225L shLUC and shRor2 cells with or without integrin α5 inhibition (10 μM ATN-191) showing cell migration within 16 h. (C) Quantification of the ratio of remaining wound area after 16 h to the initial wound area with or without integrin α5 inhibition. (D) Western blot of 2225L shLUC and shRor2 organoids for phosphorylated FAK and total FAK after 10 μM ATN-191 treatment. (E) Western blot of 2225L shLUC and shRor2 cells for phosphorylated FAK and total FAK after 0, 10, 50, 100, or 200 nM PF-562271 Besylate treatment. (F) Brightfield images of wound healing assays on 2225L shLUC and shRor2 cells with or without FAK inhibition (200 nM PF-562271 Besylate) showing cell migration within 16 h. (G) Quantification of the ratio of remaining wound area after 16 h to the initial wound area with or without FAK inhibition. (H) Western blot of 2225L shLUC and shRor2 organoids for phosphorylated FAK and total FAK after 200 nM PF-562271 Besylate treatment. (I) Brightfield DIC images showing compromised invasion of shRor2 organoids into the surrounding matrix after integrin α5 or FAK inhibition. Scale 30 μm. (J) Quantitation of cellular projections emanating into the surrounding matrix in shRor2 vs. shLUC organoids (p-value noted in the panel; ns: not significant; n=23, 9, 16, 22, 10, and 16 organoids for shLUC+vehicle, shLUC+integrinα5 inhibition, shLUC+FAK inhibition, shRor2+vehicle, shRor2+integrinα5 inhibition, and shRor2+FAK inhibition group, respectively).

## DISCUSSION

Cancer cells exhibit extremely adaptable cellular programs enabling their successful dissemination, survival, transit, and establishment of distant metastases. Such proficiency in migration and invasion mechanisms requires extensive cell-cell and cell-ECM exchanges along with molecular signaling events which guide and mobilize tumor cells. We demonstrate that the noncanonical Wnt receptor, Ror2, regulates intercellular adhesion and cell-ECM interactions impacting tumor cell invasion and ECM composition to facilitate cancer cell invasion. We discovered that compromised Wnt/Ror2 signaling *in vivo* and within 3D-cultured tumor organoids specifically disrupts actin dynamics, adhesion, and tumor cell-intrinsic ECM deposition, including collagen crosslinking gene expression programs likely conducive for reinforcing tumor cell transit from the primary tumor. Interestingly, E-cadherin downregulation was observed upon Ror2 loss, particularly at invading tumor cell protrusions within the surrounding ECM (**Figure 1**). Integrin receptors, specifically α5 integrin and β3 integrin, were also upregulated and spatially repositioned to the invasive front and cell-matrix interface of Ror2-deficient tumor cells and organoids (**Figure 2**), similar to roles for integrins directional migration maintenance in prostate cancer cells^30^ and fibroblasts^31,32^, as well as developmental processes encompassing convergent extension and oriented cell division^33,34^. This change in integrin presence and clustering was also accompanied by the simultaneous production of a provisional fibronectin matrix, a vital component of the ECM, ligand for α5 integrin, and mediator of collagen assembly and organization^35^. Along with altered ECM architecture, Ror2 downregulation changed the intratumor spatial landscape of integrin and FAK activation within primary tumors (**Figures 2, 3**), suggesting an important physiological function for Ror2 in shaping both signaling and ECM architecture during tumor progression. Blocking either integrin or FAK, a downstream mediator of integrin-mediated signal transduction, inhibited the migration and dissemination observed upon Ror2 depletion (**Figure 5**).

The precise cellular and molecular mechanisms regulating tumor cell invasion and dissemination during metastasis remain to be elucidated. The epithelial to mesenchymal transition (EMT) has gained considerable attention as a mediator of cancer cell migration, particularly through the binary loss of adherens junction components (E-cadherin) and tight junctions (claudins) and upregulation of mesenchymal transcription factors Snail, Twist, and Zeb1^36,37^. However, emerging evidence now demonstrates that collective cell migration likely also contributes to invasion and metastasis in cancer^38^. Like normal developmental contexts where the biological process is building a tissue or healing a wound, tumors also encompass cell-rich environments that range in form and function^13^. Detailed analysis of the EMT process has helped define mechanisms responsible for cancer progression, particularly aspects of cellular plasticity, treatment resistance, and intermediary cell states during cancer cell adaptation and tumor evolution^37,39,40^. Nonetheless, defining the regulation of specific cell-cell and cell-ECM interactions during tumor progression is a necessary step to understand the interplay of cell signals guiding multicellular tumor cell behavior. In the present study, we establish that depletion of Ror2 levels in tumor cells disrupts the levels and localization of E-cadherin within tumor cell junctions. Although we occasionally observed the concomitant expression of the mesenchymal marker, vimentin, in invasive areas associated with E-cadherin loss (**Figure 1**), we did not observe the overt upregulation of mesenchymal genes in shRor2 organoids, suggesting the downregulation of junctional E-cadherin predominantly occurred in the absence of a distinct mesenchymal switch after Ror2 downregulation. This observation has important implications in cancer cell invasion and metastatic progression given that fluctuations in E-cadherin levels, dictated by Ror2, might play an essential role in dictating states of cell adhesion, dissemination, and survival during metastatic progression^41,42^. Importantly, E-cadherin levels are diminished but not completely lost upon Wnt/Ror2 disruption. This type of hybrid EMT state, rather than complete mesenchymal state, is associated with tumor cell aggression in metastasis^43-45^. Beyond E-cadherin as a marker of the epithelial state in EMT, roles for E-cadherin include the regulation of epithelial organization and polarity^46^, as well as the modulation of growth factor signals like EGF^47,48^. In line with such roles for E-cadherin, the regulation of junctional mechanocoupling by Wnt5a/Ror2 has been observed during angiogenic collective migration, where Wnt5a activated Cdc42 at cell junctions and stabilized vinculin/α-catenin binding to support adherens junction coupling with the actin cytoskeleton^49^. Interestingly, vinculin levels were upregulated as a consequence of Ror2 deletion in our models, suggesting potential compensatory mechanisms in response to E-cadherin loss and altered mechano-regulation of cell-cell adhesion in tumor cells^50^.

The alteration in tumor stroma, particularly the increased presence of Type I collagen upon Wnt/Ror2 disruption (**Figure 1**), prompted our investigation of Wnt/Ror2 signaling in controlling cell-cell and cell-matrix interactions in TNBC. Increased ECM stiffness within the tumor microenvironment is associated with poor patient prognosis across breast and other cancer types^9,51,52^. Enhanced cross-linked collagen within the ECM milieu contributes to ECM stiffness, promoting tumor cell invasion and metastasis^53^. Our data now suggest that ECM composition and tumor cell phenotype are shaped by cell intrinsic spatiotemporal Wnt modes of signaling within the tumor landscape. In addition to the reinforcement of collagen crosslinkers within the lysyl oxidase gene family (lox, loxl2, **Figure 1**), the heightened integrin expression upon Ror2 loss, particularly α5 and β3 integrins, prompted us to investigate the composition and reinforcement of ECM cues that enable Ror2-deficient tumor cells to migrate and invade. Fibronectin is a known structural ECM protein which binds α5 integrin, and its polymerization is known to regulate the composition and stability of extracellular matrix fibrils, particularly type I collagen^54,55^. Intriguingly, we not only detected the clustering and upregulation of α5 integrin at sites of tumor cell invasion, but we also detected the deposition of fibronectin specifically at the interface of Ror2-depleted tumor cells with pericellular ECM (**Figure 2**). Interestingly, while dnTcf4-mediated inhibition of β-catenin activation did not impact tumor cell invasion prompted by Ror2 loss, β-catenin-dependent signaling in the absence of Ror2 was necessary for the upregulation of *Itga5* and *Itgb3* (**Figure 4**). Moreover, β-catenin-dependent effects on tumor cells were distinct from Wnt/Ror2 disruption with respect to collagen crosslinking, matrix deposition and composition, and actin cytoskeleton changes, suggesting that Wnt/β-catenin-dependent and Wnt/Ror2 signaling differentially modulate ECM ligand-receptor repertoires to shape tumor cell behaviors. Wnt signaling has been implicated in the setting of fibrosis and wound healing^56-58^, yet it remains unclear how spatiotemporal control of cell-ECM interactions is achieved by distinct, but integrated, modes of Wnt signaling across a multicellular context. Our studies have several implications on tumor progression, specifically where the integration of Wnt signaling modes is present among subpopulations within the cell-rich landscape of TNBC^15^.

Our results may help explain how fibronectin expression, guided by Ror2 presence, provides key signals for tumor cells to survive, directionally orient, and move through the local microenvironment to invade and eventually disseminate from the primary tumor^31^. Key studies in *Xenopus* demonstrate that integrin-fibronectin interactions can specifically polarize actin-rich protrusions, and such protrusive activity exerted by migrating cells subsequently help to fuel polymerization and orientation of ECM fibrils through tractional forces generated by the migrating cell^59-61^. Moreover, given that a polymerized fibronectin network is also important for the assembly of other ECM-associated constituents like collagens^54^, heparin sulfate proteoglycans^62^, and tenacin C^63^, our results place Wnt/Ror2 signaling as an important nexus for tumor cell exchanges with the ECM. Ror2 downmodulation *in vivo* drastically augmented integrin, collagen, and fibronectin presence within tumor cells and changed the topology of cell-ECM signaling *in vivo* such that focal adhesion activation was expanded in Ror2-depleted vs. Ror2-intact tumors (**Figures 1-3**). Interestingly, matrix rigidity has been shown to impact the amplitude of Wnt/β-catenin activity through downmodulation of the negative regulator Dikkopf-1 (Dkk1)^64^. Whether such feedback mechanisms apply in our studies remains an open question. Integrins are critical ECM receptors which relay extracellular signals to receiving cells, and their content and activity can dictate reciprocal cell-ECM responses during malignancy^9^. The fact that integrins, fibronectin, and other ECM components, identified downstream of Wnt/Ror2 signaling in this study, have been implicated in the regulation of tumor cell dormancy and survival during cancer metastasis^65-67^ warrants further investigation. These data, together with other studies that demonstrate fibronectin can promote directional persistence of cancer cell invasion^68^, suggest that alternative Wnt/Ror2 signaling orchestrates critical tumor cell-ECM exchanges within TNBC by elaborating matrix composition and specific interactions between the cancer cell and its microenvironment.

Aside from Wnt/β-catenin signaling, Wnt/PCP signaling through Wnt/Glipican4/Frizzled has been shown to regulate ECM assembly through effects on cadherin-mediated cell cohesion^33^. VANGL/Prickl1a have opposing functions on ECM organization with respect to fibronectin assembly in zebrafish embryos. Other studies have implicated Wnt/PCP as an essential mediator of integrin transmission of cytoskeletal tension required to direct fibronectin fibril formation at cell surfaces during convergent extension in Xenopus embryos^69^. The downregulation of cofilin and shift in F-actin/G-actin ratios upon Ror2 loss suggest that Wnt/Ror2 signaling is an important determinant for tumor cell-intrinsic ECM production and response by modulating actin polymerization and depolymerization. Roles for cofilin in migration and turning of metastatic cancer cells exist, highlighting the rather intricate functions such proteins perform in directing deliberate tumor cell movements^70^. Known roles also exist for Wnt signaling in reorganizing the actin cytoskeleton in various developmental and cancer contexts across metazoans, particularly though Rho GTPase interactions^71,72^. Interestingly, during zebrafish gastrulation, Wnt11 controls tissue morphogenesis by modulating E-cadherin-mediated cell cohesion through Rab5c-dependent actin remodeling^73^.

Based on these observations, critical questions remain of how the spatial integration and interplay of Wnt pathways regulate cellular diversity and tumor cell behavior during cancer progression. Given our previous findings that canonical and noncanonical Wnt signaling modes are integrated within mammary development and breast cancer ^15,28^, the characterization of Wnt/Ror2 and other Wnt signaling modes in shaping the evolutionary landscape of tumor cells during breast cancer metastasis will be of interest. Defining such signaling pathways that shape adhesive, migratory, and survival states within a tumor will provide a more comprehensive understanding of the phenotypic variation of tumor cells and the spatiotemporal states of signaling guiding both the composition and cooperativity of cell-ECM interactions during breast cancer progression.

## Materials and Methods

### Mouse strains

This study is compliant with the rules of the Guide for the Care and Use of Laboratory Animals of the National Institutes of Health. The animal maintenance and procedures are approved by the Baylor College of Medicine Institutional Animal Care and Use Committee (Protocol AN-504). A transplantable TP53-null mammary tumor bank was generated as described ^21^. Basal-like 2225L and 2153L models, maintained as a transplantable biobank, were propagated in the Balb/c inbred female mice (strain #047, 3–4 weeks of age, ENVIGO Houston, TX, USA).

### Tumor propagation, cell isolation, and cell culture maintenance

To generate shLUC and shRor2 2225L tumors, primary tumor cells were isolated and transduced as previously described^15^. Cells were then injected into cleared mammary fat pads at 25,000 cells/10 μl injection volume. After 3-4 weeks of propagation, mammary tumors were monitored by caliper measurement and collected at a volume of approximately 500mm^3^. Tumor tissues were minced into 1×1 mm fragments and enzymatically dissociated in HBSS containing 1 mg/ml collagenase A (#11088793001; Roche, Basel, Switzerland) and 1 μg/ml DNase I (#07900; Stemcell Technologies, Vancouver, Canada) in a shaking incubator at 37 °C, rotating at 125rpm for 2 h. Tissue digests were pipetted and mixed every 20 min to facilitate homogeneous dissociation. To enrich for tumor organoids, digests were subjected to differential centrifugation at 1500rpm for 10 seconds, repeating 3 times. Enriched organoids were further dissociated using TrypLE at 37 °C for 5 min and washed and filtered through a 40 μm cell strainer to obtain single cells. Single cells were washed with 1X DPBS and resuspended in appropriate media further experiments.

For 2D culture, single cells were cultured in DMEM/F-12 medium (#11320033; ThermoFisher Scientific, Waltham, MA, USA) supplied with 10% FBS (#10082147; ThermoFisher Scientific), 5 μg/ml insulin (#I5500; Sigma-Aldrich, St. Louis, MO, USA), 1 μg/ml hydrocortisone (#H0888; Sigma-Aldrich), and 10 ng/ml mEGF (#SRP3196, Sigma-Aldrich). 200 µg/ml Geneticin (#10131035; ThermoFisher Scientific) was added to select for tumor cells. For organoids culture, single cells (500,000 cells/well) were seeded in Ultra-low attachment 24-well plates (#3473, Corning, Corning, NY, USA) to enable the formation of cell aggregates. Aggregated organoids were then used in tumor organoids assays.

### Generation and transduction of lentiviral shRNAs

Lentiviral LeGO plasmids encoding shLUC and Ror2 shRNA hairpin sequences were derived and validated in our prior studies ^15,28^. shRor2-1 and shRor2-2 hairpin sequences correspond with shRor2-94 and -98 clones from the MISSION pLKO Lentiviral shRNA libraries (Sigma). Wnt pathway reporters (plasmid 7TGC #24304, Addgene, Cambridge, MA, USA) were validated in mammary epithelial cells as described^15,28^. For dnTcf4 (pCWXPGR-pTF-dnTCF4, Plasmid #114277) and β-catenin (pCWXPGR-pTF-betaCatenin, Plasmid #114281) lentiviral studies, plasmids were a gift from Patrick Salmon and acquired from Addgene. Tumor cells (500,000 cells/well) were infected in 24-well low-attachment plates or monolayer culture at an MOI of 30 with lentivirus in growth media. Cells were briefly rinsed three times with phosphate-buffered saline after overnight infection, and aggregated organoids or monolayer cells were then used in downstream experiments.

### Tumor organoids assays

The organoids assays were performed in 2 mg/ml Type I collagen matrices. Rat-tail collagen Type I (#354236; Corning) was diluted in 10x MEM, sterile dH2O, and 7.5% sodium bicarbonate solution to achieve a neutralized 2 mg/ml collagen solution at pH 7.0. Eight-well chamber slides were coated with 5 μl collagen and incubated at 37 °C for 15 min. Aggregated organoids were washed in phosphate-buffered saline before being suspended in collagen at 50,000 cells/40μl volume. 40 μl of cell suspension was plated into each chamber and the chamber slides were incubated at 37 °C for 1 hr. After polymerization, collagen gels were overlaid with 500 μl of growth media. For RNA and protein extractions, collagen matrices were dissociated in 2 mg/ml collagenase A solution at 37°C for 20 minutes under constant rotation at 40rpm. Organoids were then washed with 1X DPBS and collected by a series of short centrifugation steps prior to cell lysis.

### RNA isolation, sequencing, and analyses

Total RNA was extracted and purified using RNeasy mini kit following the manufacturer’s protocol (#74104; Qiagen, Germantown, MD, USA). The BCM Genomic and RNA Profiling Core performed sample quality checks using the Nanodrop ND-1000 (ThermoFisher Scientific) and a Bioanalyzer Nano chip (Agilent Technologies, Santa Clara, CA, USA). RNA integrity and RNA-seq library preparation were conducted by the Genomics and RNA sequencing Core, followed by Next Gen Sequencing on the NovaSeq 6000 Illumina platform. Following acquisition, post-sequencing analysis was conducted by performing sequencing alignment and transcript abundance using STAR or HISAT2 and Cufflinks. Fragment per kilobase of transcript per million mapped reads (Fpkm) values were log2-transformed prior to analysis, where data were compared to analyze gene expression changes between shLUC and shRor2 groups. Genes were identified as significantly altered in shRor2 conditions based on p<0.01 (t-test) and fold change cutoff. The Database for Annotation, Visualization and Integrated Discovery (DAVID) was used to help to prioritize gene sets by identifying enriched biological themes, functional-related gene groups, and interacting proteins that were differentially expressed^74^. RNA sequencing data were deposited in NCBI’s Gene Expression Omnibus (GEO) under accession numbers GSE174506 (2225L shLUC vs. shRor2) and GSE176041 (2153L shLUC vs. shRor2) for GEM organoid studies.

### Quantitative real-time PCR

Total RNA was reverse transcribed using high-capacity RNA-to-cDNA kit (#4388950; Applied Biosystems, Foster City, CA, USA). Quantitative real-time PCR was performed by StepOnePlus Real-Time PCR system using SYBR Green PCR master mix (Applied Biosystems). Primer sequences (Supplementary Table S1) were designed using the NCBI nucleotide BLAST. GAPDH was used as a reference gene for normalization, and relative gene expression fold changes were calculated as 2^-ΔΔCt^.

### Processing of tumors tissue and organoids cultures

5-Bromo-2’-deoxyuridine (Brdu) at 60 μg/g body weight was injected into the mice via intraperitoneal injection 2 h before tissue collection. Tumors were dissected and fixed in 4% paraformaldehyde overnight at 4 °C before processing to paraffin blocks. Organoid cultures were washed with phosphate-buffered saline and fixed in 4% paraformaldehyde for 5 min at room temperature before processing to paraffin blocks. Prior to processing in paraffin block, organoids in 3D collagen were oriented in HistoGel (Epredia™, #22-11-678) to help maintain orientation and integrity prior to processing. Paraffin-embedded tumor tissue or 3D organoids were sectioned at 5μM thickness prior to immunostaining.

### Immunostaining

Tissue and organoid sections were deparaffinized, rehydrated and subjected to heat-induced epitope retrieval using sodium citrate pH 6.0 or Tris-EDTA pH 9.0 antigen retrieval for 20 min. Sections were blocked at room temperature for 1 h by MOM blocking solution (#BMK-2202; Vector Laboratories, Burlingame, California, USA) containing 5% BSA. Primary antibodies were applied overnight at 4 °C. Antibodies and concentrations were: Ror2 (1:500; Developmental Studies Hybridoma Bank, Iowa City, IA, USA), eGFP (1:1000; #ab290; Abcam, Cambridge, MA, USA), RFP (1:1000; #600-401-379; Rockland, Pottstown, PA, USA), Integrin α5 (1:1000; #ab150361, Abcam), Fibronectin 1 (1:2000; #610077; BD Biosciences, San Jose, CA, USA), phospho-FAK (1:1000; #44-626G; ThermoFisher Scientific), phospho-ERM (1:1000; CST, Danvers, MA, USA), cofilin (1:1000; #5175S; CST), and phospho-cofilin (1:500; #3313S; CST). Sections were washed three times in 1X PBS and incubated with Alexa Fluor 488-conjugated or Alexa Fluor 594-conjugated goat anti-rabbit or anti-mouse IgG secondary antibodies in MOM diluent containing 5% BSA in the dark for 1 h at room temperature. Sections were washed three times in 1X PBS and counterstained with 1 μg/ml DAPI before mounting slides with ProLong Diamond Antifade Mounting Media (P36961; ThermoFisher Scientific). Tyramide amplification was performed for Ror2 and phospho-FAK detection according to the manufacturer’s instructions (#NEL701A001KT; PerkinElmer, Waltham, MA, USA).

### Western blotting

Protein samples were separated in NuPAGE 4-12% Bis-Tris protein gels (#NP0336BOX; ThermoFisher Scientific) and transferred to PVDF membrane (#LC2002; ThermoFisher Scientific) for antibody probing. The blots were blocked with 5% blocker (#1706406; Bio-Rad, Hercules, CA, USA) in Tris-buffer saline containing 0.05% Tween-20. Blots were then incubated with primary antibodies overnight at 4 °C before incubating with HRP-conjugated secondary antibodies for 1 h at room temperature. The following antibodies were used in western blotting: Ror2 (1:1000, Developmental Studies Hybridoma Bank), Integrin α5 (1:1000; #ab150361, Abcam), Fibronectin 1 (1:1000; #610077; BD Biosciences), FAK (1:1000; #3285; CST), phospho-FAK (1:1000; #44-626G; ThermoFisher Scientific), cofilin 1 (1:1000; #5175S; CST), and GAPDH (1:2500; #5174; CST).

### F-actin/G-actin assay

Approximately 1 × 10^7^ shLUC and shRor2 cells were harvested. F-actin and G-actin protein samples were collected according to the manufacturer’s instructions (#BK037, Cytoskeleton Inc., Denver, CO, USA). F-actin and G-actin were quantified by SDS-PAGE and western blot using actin antibody (#BK037, Cytoskeleton Inc.).

### Microscope image acquisition and analysis

Confocal imaging and IncuCyte live imaging for this project were carried out within the Integrated Microscopy Core (IMC) and Optical Imaging and Vital Microscopy Core (OiVM) at Baylor College of Medicine. In the IMC, confocal images were acquired with a Nikon A1-Rs confocal microscope with 20x (air), 40x (air), 40x (oil), and 100x (oil) objectives. Color channels were merged and converted to RBG images using Fiji 1.53c. Live imaging of 2D wound healing was performed on an IncuCyte S3 system. Single-plane or confocal stacks were analyzed using Fiji 1.53c. Super-resolution confocal imaging was performed within the OiVM Core using a Leica LSM 880 with Airyscan FAST microscope equipped with a 34-channel spectral array with laser lines at 405nm, 488nm, 514nm, 561nm, 594nm, and 633nm. Leica LSM 880 images were captures with 10x (air), 20x (air), or 40x (oil) objectives. Post-processing and image analyses were performed in Fiji 1.53c. For measurements of collagen integrated density, color-based thresholding was used to segment the blue regions, after the Analyze tool was used to measure the integrated density of the blue collagen regions within each tumor. Integrated density is Area (number pixels) x Total Intensity of pixels. For fluorescence intensity calculations, multiple regions of known area were selected within tumors using the circular draw tool in Fiji. Within the Analyze menu, Area integrated intensity and mean grey values were selected for measurement. Background fluorescence was acquired from an area within the specimen lacking fluorescence and this value was subtracted to arrive at the corrected mean fluorescence intensity.

### Wound healing assay

Cells (50,000 cells/well) were seeded into 2-well culture inserts (ibidi, Martinsried, Planegg, Germany) in 12-well plates. Medium containing 1% FBS (200μl) and indicated DMSO/inhibitors were added to each insert well. Inserts were removed on the following day, and 1 ml of fresh medium containing 1% FBS and indicated DMSO/inhibitors were added into each well. Live imaging was performed using an IncuCyte S3 system and the images were analyzed using Wound Healing Tools in Fiji 1.53c^75^. The time-lapse movies were generated by Fiji 1.53c.

### Statistical analysis and rigor and reproducibility

Data are expressed as the mean ± standard deviation (sd); n represents the number of biological replicates, unless specifically indicated otherwise in the figure legend. One-way ANOVA with Tukey’s multiple comparison tests were performed on multi-group comparisons. Unpaired two-tailed Student’s t-tests were performed on analyses involving two-group comparisons unless otherwise noted. Quantitative measurements were performed in ImageJ or GraphPad Prism 9. P<0.05 was considered statistically significant in all analyses, where ^*^ P<0.05,; ^**^ P<0.01, ^***^P<0.001, ^****^ P<0.0001). All experiments were reproduced across multiple (≥3) biological replicates.

## Figure Legends

**Figure S1**. (A) Heatmap display of significantly differentially expressed genes (p<0.01 and fold change>1.4) in 2153L shLUC and shRor2 organoids by RNA sequencing. Fold changes are represented by two-way gradients to blue (downregulation) and orange (upregulation). (B) Gene ontology analysis demonstrating the enrichment of gene expression in particular biological processes in 2153L shRor2 organoids. (C) Flow cytometry analysis of Itgb3 (CD61) in shLUC vs. shRor2 2225L tumor cells.

**Figure S2**. (A) Western blot for Fn1 and Cf1 levels following 72 h induction of dnTcf4 in 2225L tumor cells. No changes were evident. (B) Western blot for Fn1 and Cf1 levels following 72 h induction of β - catenin in 2225L tumor cells. No changes were evident. (C) RT-qPCR for mesenchymal markers *Vim, Zeb1, Snai1, Snai2* after inducible activation of Wnt/β-catenin following 72 h induction by doxycycline administration. Upregulation of EMT-associated genes is shown for *Vim, Zeb1, and Snai1*.

## Acknowledgements

The authors declare no competing financial interests. KR is supported by NCI Transition Career Development Award 5K22CA207463, Susan G. Komen CCR18548284, a Dan L. Duncan Comprehensive Cancer Center Faculty Scholar Award, L.E Gordon Cancer Research Fund, Caroline Wiess Law Fund for Research in Molecular Medicine, and NCI Breast SPORE Career Enhancement Award 2P50CA186784. HS and NZ were additionally supported by CA016303-45 and CA148761-11, respectively. We also thank Fengju Chen for bioinformatics support within the Biostatistics and Cancer Bioinformatics Division of the DLDCCC. This project was also supported by the Genomic and RNA Profiling Core at Baylor College of Medicine and the Cytometry and Cell Sorting Core at Baylor College of Medicine with funding from the CPRIT Core Facility Support Award CPRIT-RP180672 and the NIH P30 CA125123 and S10 RR024574.

## Notes

### Competing Interest Statement

The authors have declared no competing interest.

### Summary of Updates

This is a revised manuscript with new data, building on our previous preprint submission.

## References

1. Chiang, A.C. & Massague, J. Molecular basis of metastasis. N Engl J Med 359, 2814–2823 (2008).

2. Friedl, P. Prespecification and plasticity: shifting mechanisms of cell migration. Curr Opin Cell Biol 16, 14–23 (2004).

3. Friedl, P. & Alexander, S. Cancer invasion and the microenvironment: plasticity and reciprocity. Cell 147, 992–1009 (2011).

4. van Amerongen, R. & Nusse, R. Towards an integrated view of Wnt signaling in development. Development 136, 3205–3214 (2009).

5. Angers, S. & Moon, R.T. Proximal events in Wnt signal transduction. Nat Rev Mol Cell Biol 10, 468–477 (2009).

6. Marusyk, A., Janiszewska, M. & Polyak, K. Intratumor Heterogeneity: The Rosetta Stone of Therapy Resistance. Cancer Cell 37, 471–484 (2020).

7. Kauppila, S., Stenback, F., Risteli, J., Jukkola, A. & Risteli, L. Aberrant type I and type III collagen gene expression in human breast cancer in vivo. J Pathol 186, 262–268 (1998).

8. Provenzano, P.P., et al. Collagen density promotes mammary tumor initiation and progression. BMC Med 6, 11 (2008).

9. Levental, K.R., et al. Matrix crosslinking forces tumor progression by enhancing integrin signaling. Cell 139, 891–906 (2009).

10. Rottenberg, S., et al. High sensitivity of BRCA1-deficient mammary tumors to the PARP inhibitor AZD2281 alone and in combination with platinum drugs. Proc Natl Acad Sci U S A 105, 17079–17084 (2008).

11. Erler, J.T., et al. Hypoxia-induced lysyl oxidase is a critical mediator of bone marrow cell recruitment to form the premetastatic niche. Cancer Cell 15, 35–44 (2009).

12. McLean, G.W., et al. The role of focal-adhesion kinase in cancer - a new therapeutic opportunity. Nat Rev Cancer 5, 505–515 (2005).

13. Friedl, P. & Gilmour, D. Collective cell migration in morphogenesis, regeneration and cancer. Nat Rev Mol Cell Biol 10, 445–457 (2009).

14. Franz, C.M., Jones, G.E. & Ridley, A.J. Cell migration in development and disease. Dev Cell 2, 153–158 (2002).

15. Roarty, K., Pfefferle, A.D., Creighton, C.J., Perou, C.M. & Rosen, J.M. Ror2-mediated alternative Wnt signaling regulates cell fate and adhesion during mammary tumor progression. Oncogene (2017).

16. Cox, T.R. & Erler, J.T. Remodeling and homeostasis of the extracellular matrix: implications for fibrotic diseases and cancer. Dis Model Mech 4, 165–178 (2011).

17. Kai, F., Drain, A.P. & Weaver, V.M. The Extracellular Matrix Modulates the Metastatic Journey. Dev Cell 49, 332–346 (2019).

18. Pickup, M.W., Mouw, J.K. & Weaver, V.M. The extracellular matrix modulates the hallmarks of cancer. EMBO Rep 15, 1243–1253 (2014).

19. Zhang, M., et al. Identification of tumor-initiating cells in a p53-null mouse model of breast cancer. Cancer Res 68, 4674–4682 (2008).

20. Herschkowitz, J.I., et al. Comparative oncogenomics identifies breast tumors enriched in functional tumor-initiating cells. Proc Natl Acad Sci U S A 109, 2778–2783 (2012).

21. Jerry, D.J., et al. A mammary-specific model demonstrates the role of the p53 tumor suppressor gene in tumor development. Oncogene 19, 1052–1058 (2000).

22. Manie, E., et al. High frequency of TP53 mutation in BRCA1 and sporadic basal-like carcinomas but not in BRCA1 luminal breast tumors. Cancer Res 69, 663–671 (2009).

23. Langerod, A., et al. TP53 mutation status and gene expression profiles are powerful prognostic markers of breast cancer. Breast Cancer Res 9, R30 (2007).

24. Hynes, R.O. Integrins: versatility, modulation, and signaling in cell adhesion. Cell 69, 11–25 (1992).

25. Pankov, R., et al. Integrin dynamics and matrix assembly: tensin-dependent translocation of alpha(5)beta(1) integrins promotes early fibronectin fibrillogenesis. J Cell Biol 148, 1075–1090 (2000).

26. Wu, C., Bauer, J.S., Juliano, R.L. & McDonald, J.A. The alpha 5 beta 1 integrin fibronectin receptor, but not the alpha 5 cytoplasmic domain, functions in an early and essential step in fibronectin matrix assembly. J Biol Chem 268, 21883–21888 (1993).

27. Kanellos, G. & Frame, M.C. Cellular functions of the ADF/cofilin family at a glance. J Cell Sci 129, 3211–3218 (2016).

28. Roarty, K., Shore, A.N., Creighton, C.J. & Rosen, J.M. Ror2 regulates branching, differentiation, and actin-cytoskeletal dynamics within the mammary epithelium. J Cell Biol 208, 351–366 (2015).

29. van Amerongen, R., Fuerer, C., Mizutani, M. & Nusse, R. Wnt5a can both activate and repress Wnt/beta-catenin signaling during mouse embryonic development. Dev Biol 369, 101–114 (2012).

30. Joshi, R., Goihberg, E., Ren, W., Pilichowska, M. & Mathew, P. Proteolytic fragments of fibronectin function as matrikines driving the chemotactic affinity of prostate cancer cells to human bone marrow mesenchymal stromal cells via the alpha5beta1 integrin. Cell Adh Migr 11, 305–315 (2017).

31. Gopal, S., et al. Fibronectin-guided migration of carcinoma collectives. Nat Commun 8, 14105 (2017).

32. Missirlis, D., Haraszti, T., Kessler, H. & Spatz, J.P. Fibronectin promotes directional persistence in fibroblast migration through interactions with both its cell-binding and heparin-binding domains. Sci Rep 7, 3711 (2017).

33. Dohn, M.R., Mundell, N.A., Sawyer, L.M., Dunlap, J.A. & Jessen, J.R. Planar cell polarity proteins differentially regulate extracellular matrix organization and assembly during zebrafish gastrulation. Dev Biol 383, 39–51 (2013).

34. Parisi, L., et al. A glance on the role of fibronectin in controlling cell response at biomaterial interface. Jpn Dent Sci Rev 56, 50–55 (2020).

35. Vega, M.E. & Schwarzbauer, J.E. Collaboration of fibronectin matrix with other extracellular signals in morphogenesis and differentiation. Curr Opin Cell Biol 42, 1–6 (2016).

36. Miettinen, P.J., Ebner, R., Lopez, A.R. & Derynck, R. TGF-beta induced transdifferentiation of mammary epithelial cells to mesenchymal cells: involvement of type I receptors. J Cell Biol 127, 2021–2036 (1994).

37. Nieto, M.A., Huang, R.Y., Jackson, R.A. & Thiery, J.P. Emt: 2016. Cell 166, 21–45 (2016).

38. Williams, E.D., Gao, D., Redfern, A. & Thompson, E.W. Controversies around epithelial-mesenchymal plasticity in cancer metastasis. Nat Rev Cancer 19, 716–732 (2019).

39. Mani, S.A., et al. The epithelial-mesenchymal transition generates cells with properties of stem cells. Cell 133, 704–715 (2008).

40. Grigore, A.D., Jolly, M.K., Jia, D., Farach-Carson, M.C. & Levine, H. Tumor Budding: The Name is EMT. Partial EMT. J Clin Med 5(2016).

41. Onder, T.T., et al. Loss of E-cadherin promotes metastasis via multiple downstream transcriptional pathways. Cancer Res 68, 3645–3654 (2008).

42. Padmanaban, V., et al. E-cadherin is required for metastasis in multiple models of breast cancer. Nature 573, 439–444 (2019).

43. Pastushenko, I., et al. Fat1 deletion promotes hybrid EMT state, tumour stemness and metastasis. Nature 589, 448–455 (2021).

44. Kroger, C., et al. Acquisition of a hybrid E/M state is essential for tumorigenicity of basal breast cancer cells. Proc Natl Acad Sci U S A 116, 7353–7362 (2019).

45. Luond, F., et al. Distinct contributions of partial and full EMT to breast cancer malignancy. Dev Cell (2021).

46. Takeichi, M. Cadherin cell adhesion receptors as a morphogenetic regulator. Science 251, 1451–1455 (1991).

47. Hoschuetzky, H., Aberle, H. & Kemler, R. Beta-catenin mediates the interaction of the cadherin-catenin complex with epidermal growth factor receptor. J Cell Biol 127, 1375–1380 (1994).

48. Qian, X., Karpova, T., Sheppard, A.M., McNally, J. & Lowy, D.R. E-cadherin-mediated adhesion inhibits ligand-dependent activation of diverse receptor tyrosine kinases. EMBO J 23, 1739–1748 (2004).

49. Carvalho, J.R., et al. Non-canonical Wnt signaling regulates junctional mechanocoupling during angiogenic collective cell migration. Elife 8(2019).

50. le Duc, Q., et al. Vinculin potentiates E-cadherin mechanosensing and is recruited to actin-anchored sites within adherens junctions in a myosin II-dependent manner. J Cell Biol 189, 1107–1115 (2010).

51. Conklin, M.W., et al. Aligned collagen is a prognostic signature for survival in human breast carcinoma. Am J Pathol 178, 1221–1232 (2011).

52. Colpaert, C.G., et al. The presence of a fibrotic focus in invasive breast carcinoma correlates with the expression of carbonic anhydrase IX and is a marker of hypoxia and poor prognosis. Breast Cancer Res Treat 81, 137–147 (2003).

53. Spill, F., Reynolds, D.S., Kamm, R.D. & Zaman, M.H. Impact of the physical microenvironment on tumor progression and metastasis. Curr Opin Biotechnol 40, 41–48 (2016).

54. Sottile, J. & Hocking, D.C. Fibronectin polymerization regulates the composition and stability of extracellular matrix fibrils and cell-matrix adhesions. Mol Biol Cell 13, 3546–3559 (2002).

55. Kadler, K.E., Hill, A. & Canty-Laird, E.G. Collagen fibrillogenesis: fibronectin, integrins, and minor collagens as organizers and nucleators. Curr Opin Cell Biol 20, 495–501 (2008).

56. Wehner, D., et al. Wnt signaling controls pro-regenerative Collagen XII in functional spinal cord regeneration in zebrafish. Nat Commun 8, 126 (2017).

57. Kumawat, K., et al. Noncanonical WNT-5A signaling regulates TGF-beta-induced extracellular matrix production by airway smooth muscle cells. FASEB J 27, 1631–1643 (2013).

58. Bielefeld, K.A., et al. Fibronectin and beta-catenin act in a regulatory loop in dermal fibroblasts to modulate cutaneous healing. J Biol Chem 286, 27687–27697 (2011).

59. Alfandari, D., Cousin, H., Gaultier, A., Hoffstrom, B.G. & DeSimone, D.W. Integrin alpha5beta1 supports the migration of Xenopus cranial neural crest on fibronectin. Dev Biol 260, 449–464 (2003).

60. Boucaut, J.C. & Darribere, T. Fibronectin in early amphibian embryos. Migrating mesodermal cells contact fibronectin established prior to gastrulation. Cell Tissue Res 234, 135–145 (1983).

61. Davidson, L.A., Dzamba, B.D., Keller, R. & Desimone, D.W. Live imaging of cell protrusive activity, and extracellular matrix assembly and remodeling during morphogenesis in the frog, Xenopus laevis. Dev Dyn 237, 2684–2692 (2008).

62. Heremans, A., De Cock, B., Cassiman, J.J., Van den Berghe, H. & David, G. The core protein of the matrix-associated heparan sulfate proteoglycan binds to fibronectin. J Biol Chem 265, 8716–8724 (1990).

63. Van Obberghen-Schilling, E., et al. Fibronectin and tenascin-C: accomplices in vascular morphogenesis during development and tumor growth. Int J Dev Biol 55, 511–525 (2011).

64. Barbolina, M.V., et al. Matrix rigidity activates Wnt signaling through down-regulation of Dickkopf-1 protein. J Biol Chem 288, 141–151 (2013).

65. Barney, L.E., et al. Tumor cell-organized fibronectin maintenance of a dormant breast cancer population. Sci Adv 6, eaaz4157 (2020).

66. Osmani, N., et al. Metastatic Tumor Cells Exploit Their Adhesion Repertoire to Counteract Shear Forces during Intravascular Arrest. Cell Rep 28, 2491–2500 e2495 (2019).

67. Ghajar, C.M., et al. The perivascular niche regulates breast tumour dormancy. Nat Cell Biol 15, 807–817 (2013).

68. Erdogan, B., et al. Cancer-associated fibroblasts promote directional cancer cell migration by aligning fibronectin. J Cell Biol 216, 3799–3816 (2017).

69. Dzamba, B.J., Jakab, K.R., Marsden, M., Schwartz, M.A. & DeSimone, D.W. Cadherin adhesion, tissue tension, and noncanonical Wnt signaling regulate fibronectin matrix organization. Dev Cell 16, 421–432 (2009).

70. Sidani, M., et al. Cofilin determines the migration behavior and turning frequency of metastatic cancer cells. J Cell Biol 179, 777–791 (2007).

71. Lai, S.L., Chien, A.J. & Moon, R.T. Wnt/Fz signaling and the cytoskeleton: potential roles in tumorigenesis. Cell Res 19, 532–545 (2009).

72. Schlessinger, K., Hall, A. & Tolwinski, N. Wnt signaling pathways meet Rho GTPases. Genes Dev 23, 265–277 (2009).

73. Ulrich, F., et al. Wnt11 functions in gastrulation by controlling cell cohesion through Rab5c and E-cadherin. Dev Cell 9, 555–564 (2005).

74. Huang da, W., Sherman, B.T. & Lempicki, R.A. Systematic and integrative analysis of large gene lists using DAVID bioinformatics resources. Nat Protoc 4, 44–57 (2009).

75. Suarez-Arnedo, A., et al. An image J plugin for the high throughput image analysis of in vitro scratch wound healing assays. PLoS One 15, e0232565 (2020).

