## Supplementary figures and images for "Noncanonical Wnt/Ror2 signaling regulates cell-matrix crosstalk to prompt directional tumor cell invasion and dissemination in breast cancer"

### Supplemental Figure 1

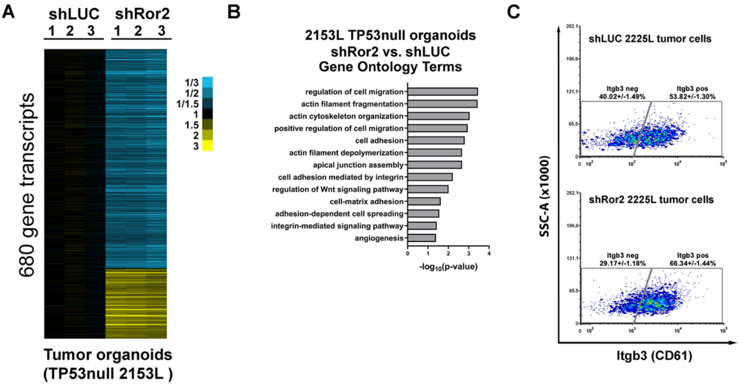

### Supplemental Figure 2

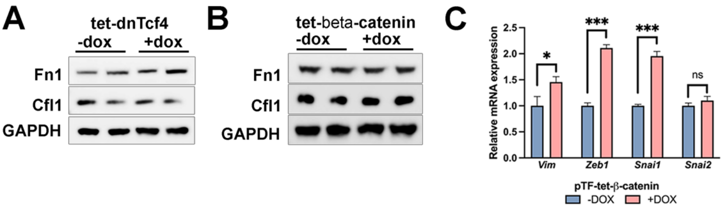
